# Design of an ultrabright biosensor for dynamic imaging of kinase activity in cells

**DOI:** 10.1101/2025.08.13.669976

**Authors:** Xiaoquan Li, Sophia K. Tan, Chan-I Chung, A. Katherine Hatstat, Qian Zhao, Jinyu Luo, William F. DeGrado, Xiaokun Shu

## Abstract

Protein kinases regulate almost every major signaling pathway. Visualizing spatiotemporal dynamics of kinase activity is thus essential to understand cell signaling. Here we report a *de <u>nov</u>o*-designed <u>a</u>ctivity reporter of <u>k</u>inase, dubbed NOVARK, which contains a single polypeptide chain with multiple modular motifs that act as specific kinase substrates and reporters. NOVARK undergoes phosphorylation-induced higher order-assembly, which are detectable as ultrabright GFP droplets with a greater dynamic range than existing Förster resonance energy transfer-based kinase reporters. We designed versions of NOVARK that rapidly and reversibly report intracellular activity of protein kinase A, C, and ERK following stimulation/inhibition by upstream GPCR agonists. Our work provides a generalizable platform that enables the design of ultrabright biosensors for illuminating dynamic architecture of kinase signaling.

---

The human genome contains over 500 protein kinases, which regulate almost all major signaling pathways by phosphorylating thousands of substrate proteins and other biomolecules (*1*-*3*). To understand role of kinases in mediating cellular signaling, it is important to visualize their dynamic activities in living cells (*4, 5*). An ideal kinase reporter should be: 1) ultrabright with a larger dynamic range than achievable with FRET; 2) temporally responsive on the same time scale as the activation/inhibition of a kinase; 3) tunable, allowing either rapid reversibility, to assess the dynamics of a kinase, or irreversibility, to evaluate a signal integrated over a time interval; 5) modular with high specificity for specific different kinases. It 6) should provide fine spatial resolution to report the location of activated kinases, and 7) it should be encoded in a single, relatively small protein that can function in a wide range of cells and cellular environments.

While recent advances in protein design and engineering have enabled the design of a number of sensors, none have addressed this entire list of features (Fig. 1A). For example, peptides and proteins that change conformation and/or aggregation state in response to phosphorylation have been described (*6*-*11*) and sensors that report phosphorylation after over 24 hours have been described (*6, 8*). Previously, we designed multi-component systems, which involve natural phosphopeptide binders, protein kinase substrates and multiple coiled coils to engineer a complex system that responds kinase activity (*12*-*14*). However, the system requires finely coordinated expression of the multiple components for each application. Here, we use concepts in *de novo* protein design and stimulus-sensitive peptide material engineering (*15*-*20*) to create sensors that fulfill the full list of desirable features. Our pipeline provides fast and ultrabright kinase reporters with large dynamic range and fine spatial resolution, which we show are useful for imaging dynamic signaling of multiple kinases including PKA and PKC following GPCR activation, and extracellular signal-regulated kinase (ERK) following receptor tyrosine kinase (RTK) activation in living cells.

**Fig. 1.**
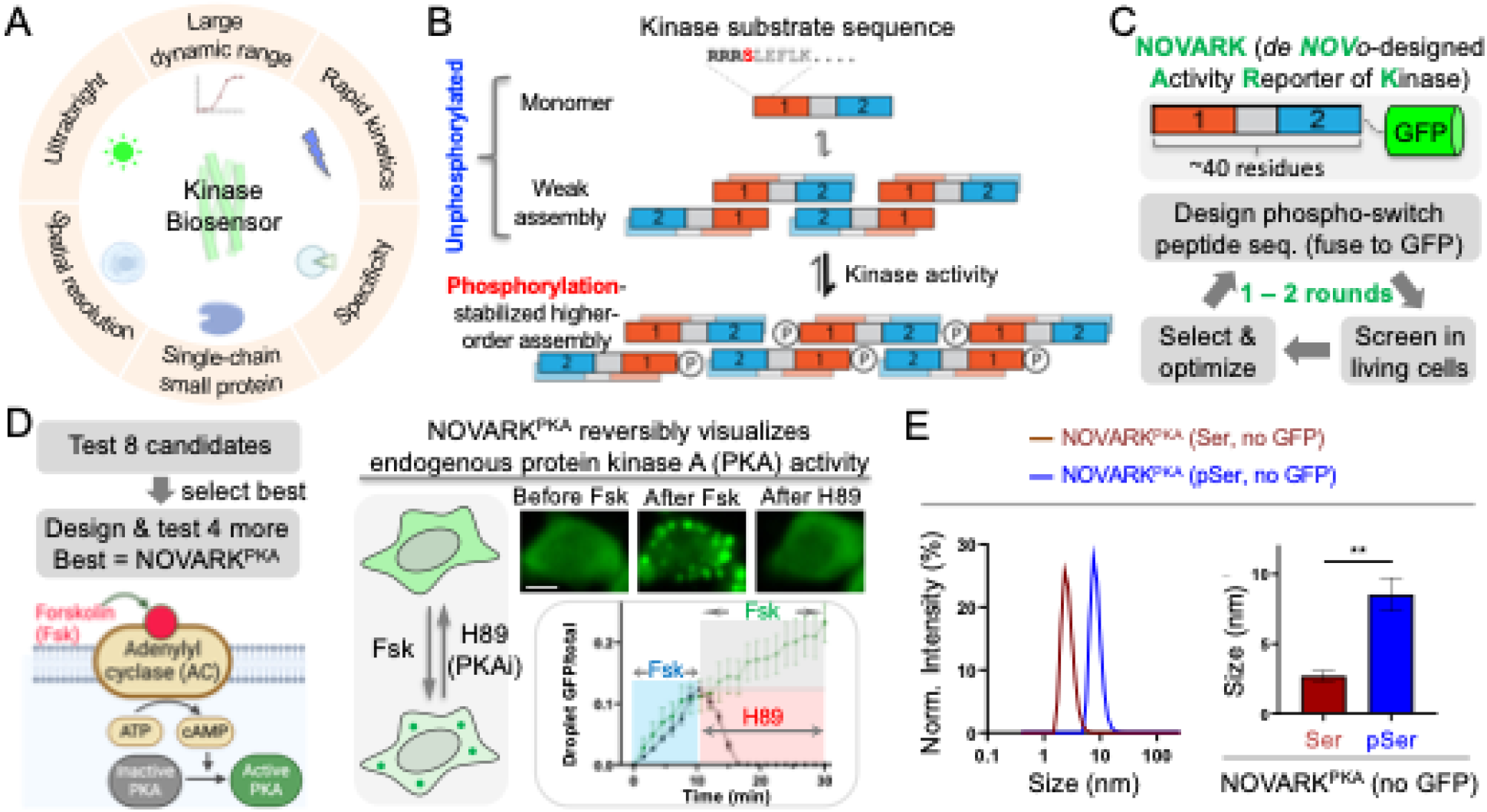
*De novo* design of phosphorylation-switch peptide-based protein kinase A biosensor. (A) Schematic showing preferred properties for an ideal kinase biosensor. (B) Schematic showing the working principles. (C) Schematic showing the engineering approach combining rational design and screening of limited number of candidates via 1 to 2 rounds. (D) Left: cartoon showing signaling pathways of forskolin-induced PKA activation. Right: fluorescence imaging of the PKA biosensor, dubbed NOVARK^PKA^. Two sets of time-lapse imaging in HEK293T cells expressing NOVARK^PKA^ were conducted: 1) Forskolin was added to the cells that were imaged for 30 minutes (the green plot); 2) Forskolin was added to the cells, and then 10 minutes later, the PKA inhibitor (PKAi) H89 was added to the cells (the black plot). Data are mean ± SE (n = 10 cells). (E) Dynamic light scattering analysis of the phosphorylation-switch peptides in NOVARK^PKA^. Data are mean ± SE (n = 3). Scale bar, 10 μm (D).

## *De novo* design of a phosphorylation-switch peptide

We designed a dynamic, self-assembling system based on coiled coils, whose stability depended on kinase-specific phosphorylation of a substrate peptide (Fig.1B). The basic module consists of a coiled-coil with staggered sticky ends to drive polymeric self-assembly. To introduce phosphorylation-dependent association, we placed kinase substrate sequences near the N-terminus of each chain in locations where phosphoserine (pSer) could interact favorably with neighboring helices in the assembly. When appropriately positioned near the N-terminus of a helix, a pSer can stabilize an α-helical conformation, shifting the equilibrium towards the assembled, helical state (*10, 21*). We also built stabilizing inter-chain interactions with the pSer residue to further promote assembly (Materials and Methods). Following computational design of the system, termed NOVARK (see Methods and Figure S1), we finalized 8 peptide sequences (table S1) based on (1) energetic and packing metrics; (2) maximization of sequence diversity between designs to cover a broad range of sequence space, and (3) fidelity of AlphaFold2(*22*) predictions to the design structure.

### Phosphorylation-switch peptide-based PKA activity reporter

To test the final 8 phosphorylation-switch peptides in mammalian cells, we genetically fused them to GFP for fluorescence imaging (Fig. 1C, table S1). These constructs were expressed in human embryonic kidney 293T (HEK293T) cells (Fig. 1D, fig. S3A – I). To activate PKA, we incubated the cells with forskolin (Fsk). Fsk binds and activates adenylyl cyclase, which then converts ATP to cyclic AMP (cAMP) that activates PKA (Fig. 1D). We found that 1 out of 8 candidates responded to PKA activation, forming higher-order assemblies (fig. S3D). The rest of the candidates showed no response (fig. S3A – C, E – H). 6 of 7 non-responsive candidates showed diffusive fluorescence before and after PKA activation (fig. S3A – C, F – H).

Interestingly, one non-responsive candidate formed fibril-like structures before PKA activation, and no further changes were observed upon PKA signaling, indicating that it forms higher-order assembly independent of phosphorylation (fig. S3E and I, movie S1). This could be useful to compartmentalize proteins of interest and manipulate their functions in living cells.

While one of the candidates showed response to PKA activity (Design D, Table S1), its response took ∼10 minutes, which is faster than previous sensors (*8*), but slower than our goal of single-digit minute time-scale – the time range of most kinase signaling response. Therefore, we decided to further optimize this candidate. We increased the dynamic range between the staggered (on-target) topology and the unstaggered (off-target) topology by placing electrostatic residues so that electrostatic interactions are maximized in the on-target conformation and penalized in the off-target conformation (*6*). This resulted in 4 new peptide sequences that were computationally predicted to confer the largest energetic gap between the on- and off-target topologies (table S1). Two of them showed faster response (one is discussed in the next paragraph, the other is discussed in the last section), and the rest showed no or slower response than the parent (fig. S4).

One optimized candidate, which contains two mutations L10E/A17K in the parent (i.e., Design D, table S1), showed dynamic response upon PKA activation. We named this peptide-based PKA reporter NOVARK^PKA^. In particular, we demonstrated that NOVARK^PKA^ showed fast response to PKA activation, forming green fluorescent punctate structures within 1.5 minutes after addition of Fsk (Fig. 1D). Furthermore, the fluorescent puncta were dissolved upon addition of the PKA inhibitor (PKAi) H89 within a timescale that is consistent with FRET-based PKA reporter (Fig. 1D) (*23*), indicating that the higher-order assembly was quickly disassembled upon PKA inhibition.

Lastly, we conducted biochemical characterization to examine whether phosphorylation of the peptide promotes larger particle formation. We synthesized the phosphorylation-switch peptide of the NOVARK^PKA^ (i.e., no GFP) with and without the phosphate group (Ser vs pSer). Circular dichroism (CD) spectroscopy revealed that both peptides formed helical structures (fig. S5). Dynamic light scattering showed that the phosphorylated peptide formed larger particles than the un-phosphorylated peptide (Fig. 1E, fig. S6). These results support our design of helical structures and phosphorylation-switch peptides.

### Generalization of the NOVARK design to PKC activity reporter

To demonstrate that our approach is generalizable to other kinases, we applied the *de novo*-designed peptide sequence (Design D) to develop a reporter for PKC. First, we substituted the PKA substrate motif (located on the first helical heptad of the peptide, Fig. 1B, table S1) with a PKC substrate sequence from previous reporters (*24, 25*). We also varied the residues located at the peptide-peptide interfaces to achieve a wide range of intermolecular affinities. 10 candidate sequences (fused to GFP) were tested in HeLa cells (table S2).

To activate PKC, we added phorbol 12-myristate 13-acetate (PMA) to the cells, which binds to the membrane and recruits and activates PKC (Fig. 2A). One candidate showed fast response within 3 minutes upon PKC activation (Fig. 2A – D), which was then named NOVARK^PKC^ (table S2). The rest showed no or slow response (fig. S7A – I): 1) 4 candidates were diffusive in the cell with no response upon PKC activation (fig. S7A, C, E, I); 2) 2 candidates formed puncta before PKC activation (fig. S7F, H); 3) 3 candidates showed slow response with ∼ 10 to 50-minute time-scale (fig. S7B, D, G).

**Fig. 2.**
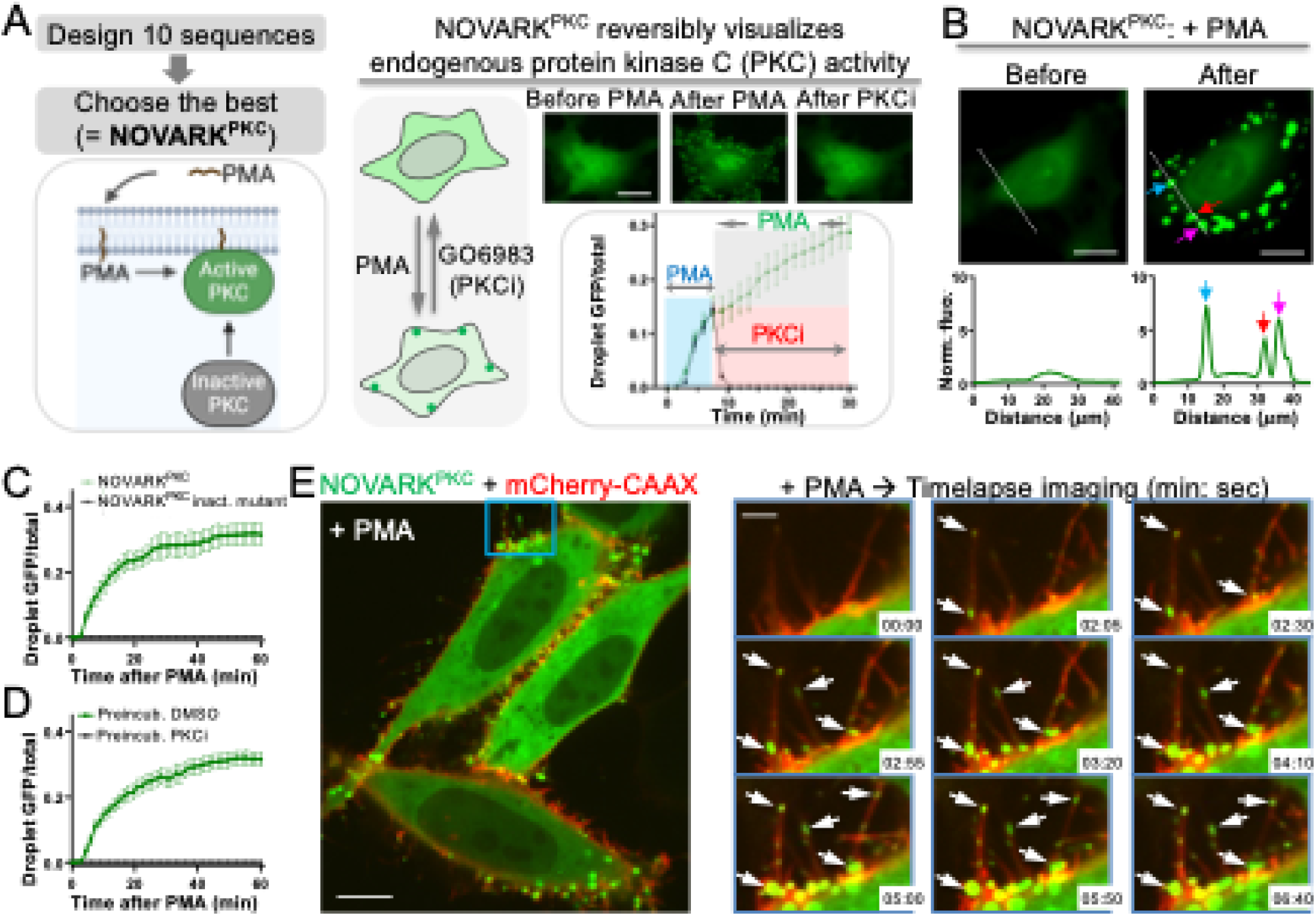
The NOVARK design is generalizable to develop protein kinase C biosensor. (A) Top-left: overview of PKC design by 1 round from 10 candidates. Bottom-left: cartoon showing signaling pathways of PMA-induced PKC activation. Right: fluorescence imaging of the PKC biosensor, dubbed NOVARK^PKC^. Two sets of time-lapse imaging in HeLa cells expressing NOVARK^PKC^ were conducted: 1) PMA was added to the cells that were imaged for 30 minutes (the green plot); 2) PMA was added to the cells, and then ∼8 minutes later, the PKC inhibitor (PKCi) GO6983 was added to the cells (the black plot). Data are mean ± SE (n = 10 cells). (B) Representative fluorescence images and histograms showing fluorescence intensity over distance along the white line indicated in the images. (C, D) Ratio of GFP in the higher-order assembly over total GFP as a function of time upon addition of PMA, for NOVARK^PKC^ and its inactive mutant that cannot be phosphorylated (C), and for NOVARK^PKC^ in cells that were pretreated with DMSO or the PKCi (D). Data are mean ± SE (n = 10 cells for C and D). (E) Multiplex imaging of HeLa cells expressing NOVARK^PKC^ and mCherry-CAAX that labels cell membrane. (Right panels) Time-lapse imaging were conducted after addition of PMA. Arrows point to the higher-order assembly of NOVARK^PKC^. The cell membrane including filopodia was shown by the red fluorescence. Scale bar, 20 μm (A&B); 10 μm (E); 3 μm (inset in E).

We then further characterized NOVARK^PKC^. First, we demonstrated that the reporter is reversible. The fluorescent puncta were dissolved upon addition of the PKC inhibitor (PKCi) GO6983 (Fig. 2A), indicating that the higher-order assembly were disassembled upon PKC inhibition. Second, we showed that puncta formation is dependent on the phosphorylation.

Mutation of this phospho-serine abolished response to PMA-induced PKC activation (Fig. 2C). Third, preincubation of the cells with the PKCi showed no PMA-induced signal (Fig. 2D).

Importantly, we demonstrated that the reporter achieved spatial resolution in visualizing PKC activity. Here we labeled the cell membrane by expressing mCherry-CAAX (*26*). Upon addition of PMA, the green fluorescent puncta formed on the plasma membrane, and even at the filopodia (Fig. 2E, movie S2). Because the membrane-located PMA recruits and activates PKC at the membrane, our data indicates that the NOVARK^PKC^ biosensor achieves spatial resolution in reporting the PKC activity.

We also demonstrated that the PKC and PKA reporters do not crosstalk. The PKC reporter NOVARK^PKC^ did not form puncta upon addition of forskolin (Fig. S8A). On the other hand, the PKA reporter NOVARK^PKA^ showed no fluorescent puncta upon addition of PMA (Fig. S8B). Therefore, our *de novo*-designed NOVARK biosensors achieve specificity in reporting the kinase signaling, and can be used as a general approach in designing reporters for most if not all kinases.

### Dynamic visualization of PKA activities during GPCR signaling

To demonstrate whether our *de novo*-designed NOVARK biosensors are applicable to biological studies, we decided to image PKA and PKC signaling that are activated through the G*α*s- and G*α*q-coupled GPCR pathway, respectively (Fig. 3A, F). For the PKA reporter NOVARK^PKA^, we applied it to visualize PKA during β2 adrenergic receptor (β2AR) and adenosine receptor signaling, both of which are coupled to G*α*s (Fig. 3A). Their agonist isoproterenol (ISO) and adenosine bind and activate β2AR and adenosine receptor, respectively. The activated G*α*s dissociates from GPCR and binds and activates adenylyl cyclase (AC), which produces cAMP. This secondary messenger then binds and activates PKA. First, we applied ISO to HEK293 cells that express endogenous β2AR-G*α*s-AC-PKA system and have been widely used for studying β2AR signaling (*27*). We showed that upon addition of ISO, NOVARK^PKA^ formed brightly fluorescent puncta within 1.5 minutes (Fig. 3B, C). Furthermore, the puncta were dissolved in ∼30 minutes, which is consistent with FRET-based PKA reporter (*28*), because ISO/β2AR induces transient PKA signaling due to breakdown of cAMP and desensitization of the receptor (*27*). Next, we demonstrated that higher-order assembly of the reporter is dependent on the phospho-serine because mutation of this residue abolished puncta formation by ISO (fig. S9A – C). Third, we showed that the reporter response is dependent on PKA activity because preincubation of the cells with PKAi abolished ISO-induced puncta formation (fig. S9D, E).

**Fig. 3.**
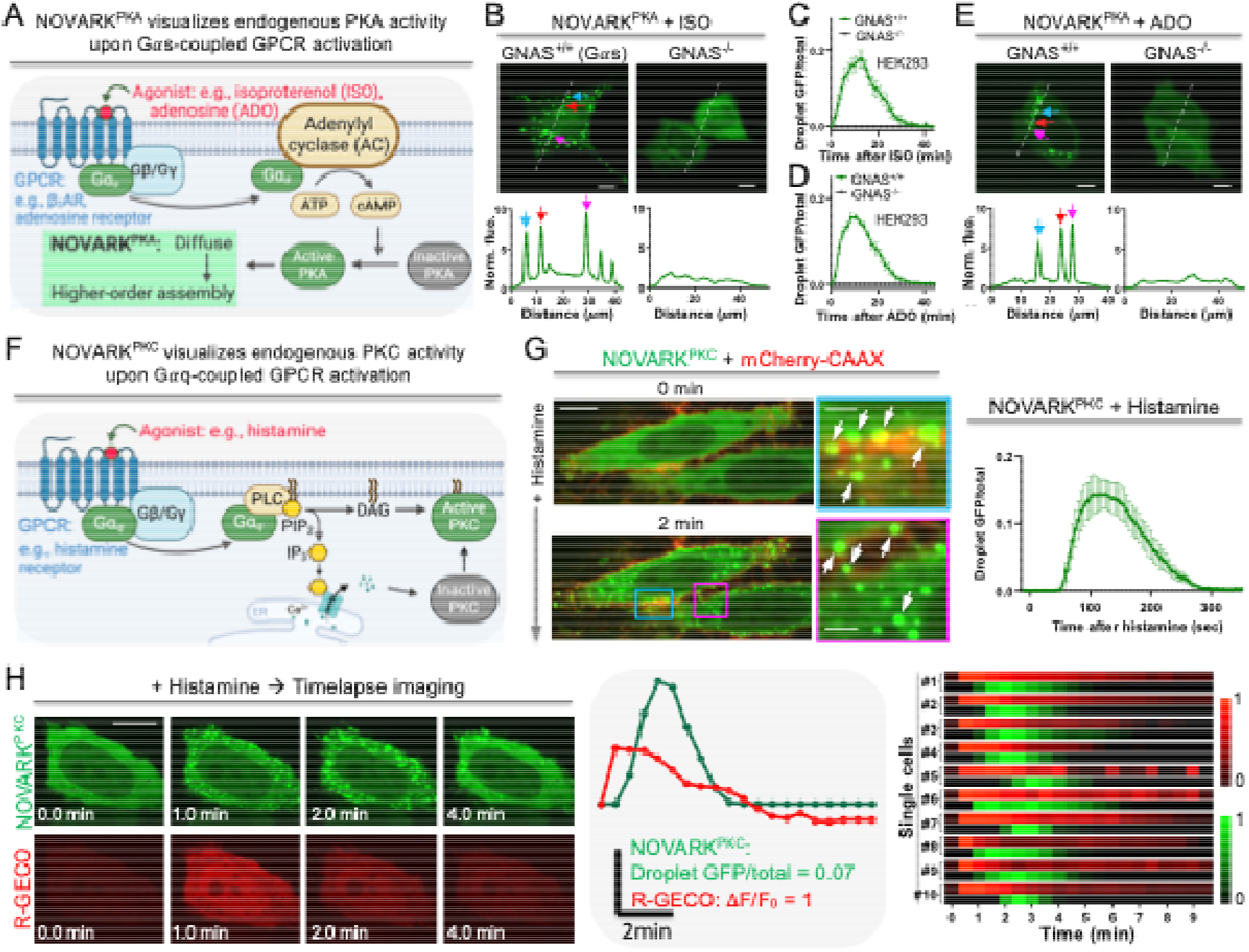
The NOVARK biosensors visualize dynamic activities of endogenous kinases during GPCR signaling. (A) Cartoon showing PKA activation in G*α*s-coupled GPCR pathway by their agonists. (B – E) Fluorescence images and quantitative analysis of parent and GNAS knock-out HEK293 cells expressing NOVARK^PKA^ upon addition of isoproterenol (B, C) or adenosine (D, E). Histograms are shown along the dotted line in the images. Data are mean ± SE (n = 10 cells for C and D). (F) Cartoon showing PKC activation in Gaq-coupled GPCR pathway by the agonist. (G) Left: multiplex imaging of HeLa cells expressing NOVARK^PKC^ and mCherry-CAAX that labels cell membrane. Arrows point to the higher-order assembly of NOVARK^PKC^. Right: ratio of GFP in the higher-order assembly over total GFP as a function of time upon addition of histamine. Data are mean ± SE (n = 10 cells). (H) Left: multiplex imaging of HeLa cells expressing NOVARK^PKC^ and the red calcium sensor R-GECO. Middle: quantitative analysis of PKC activity and calcium signaling over time after addition of histamine for the cell shown on the left. Right: the normalized kinase activity (green) and normalized calcium signaling (red) are plotted over time after addition of histamine in single cells (#1 to 10). Scale bar, 10 μm (B, E, G, H); 2 μm (inset in G).

Fourth, we applied ISO to the HEK293 cells with *GNAS* knockout (*GNAS*^-/-^) that would break the β2AR-G*α*s-AC-PKA pathway (*29*), and indeed the fluorescent puncta were no longer formed (Fig. 3B, C). These results indicate that the reporter visualizes PKA activity via the β2AR-G*α*s-AC-PKA pathway, and can be used to study β2AR-PKA signaling.

In addition to the β2AR-G*α*s-AC-PKA pathway, we further demonstrated the PKA reporter NOVARK^PKA^ in imaging PKA signaling via another G*α*s-coupled GPCR, adenosine receptor (*30*). First, we applied adenosine (ADO) to HEK293 cells that are known to express endogenous adenosine receptor. Fluorescence imaging revealed that upon addition of ADO, the PKA reporter formed green fluorescent puncta within 1.5 minutes (Fig. 3D, E). And *GNAS* knockout blocked the reporter response to ADO (Fig. 3D, E). Together, our results demonstrate that the *de novo*-designed PKA reporter NOVARK^PKA^ is able to detect endogenous PKA activity and can be used for biological study of the GPCR-PKA signaling.

### Imaging endogenous PKC activity during histamine receptor signaling

For the PKC reporter NOVARK^PKC^, we applied it to visualize endogenous PKC activity during histamine receptor signaling, which is coupled to G*α*q (Fig. 3F) (*31*). The histamine agonist binds and activates the receptor, leading to G*α*q dissociation from the GPCR. The activated G*α*q then binds and activates phospholipase C (PLC), producing diacylglycerol (DAG) and inositol triphosphate (IP3). IP3 then releases Ca^2+^ from ER into the cytosol. PKC is then activated by DAG and Ca^2+^ (Fig. 3F) (*32*). First, we added histamine to HeLa cells that express endogenous histamine receptor (*31*). We showed that upon addition of histamine, NOVARK^PKC^ formed brightly fluorescent puncta within 1 minute (Fig. 3G, movie S3). Furthermore, the puncta were dissolved in about 6 minutes, suggesting that the biosensor reports transient activation of PKC. Next, we did multiplex imaging for both PKC and Ca^2+^ using R-GECO (*33*). Our data showed that Ca^2+^ signal preceded PKC activity as expected (Fig. 3H), which is consistent with previous studies (*24, 25*).

### NOVARK-based irreversible PKA and PKC reporters

While a reversible reporter can read out both activation and inactivation of kinase activities using time-lapse imaging, an irreversible reporter will also be useful to read out historical activation of kinase signaling, which will be useful for biological studies and high throughput screening of kinase inhibitors using snapshot imaging. Therefore, we decided to develop an irreversible *de <u>nov</u>o*-designed <u>a</u>ctivity reporter of <u>k</u>inase, dubbed iNOVARK.

For the irreversible PKA reporter, we found that one of the 4 mutants that were designed based on the Design D showed sustained signal (i.e., green fluorescent higher-order assembly) after 1 hour of ISO treatment (Fig. 4A). We named this mutant iNOVARK^PKA^ (table S1). This is in contrast to the transient signal from the reversible reporter NOVARK^PKA^ that lasted for half an hour (Fig. 3B, C). GNAS knockout abolished the reporter’s response, indicating that the higher-order assembly is dependent on the *β*_2_AR signaling (Fig. 4B, C). We further showed that iNOVARK^PKA^ can also report ADO-induced PKA activity, and the fluorescent signal is also sustainable for 1 hour (Fig. 4D, E), in contrast to the transient signal by the reversible reporter (Fig. 3D, E). GNAS knockout also abolished ADO-induced reporter signal. Interestingly, the irreversible reporter formed elongated fibril-like structures (Fig. 4F), in contrast to the round droplets of the reversible reporter (Fig. 4B, E). Furthermore, the short fibrils could fuse together to form ∼10*µ*m long fibril-like structures (Fig. 4F).

**Fig. 4.**
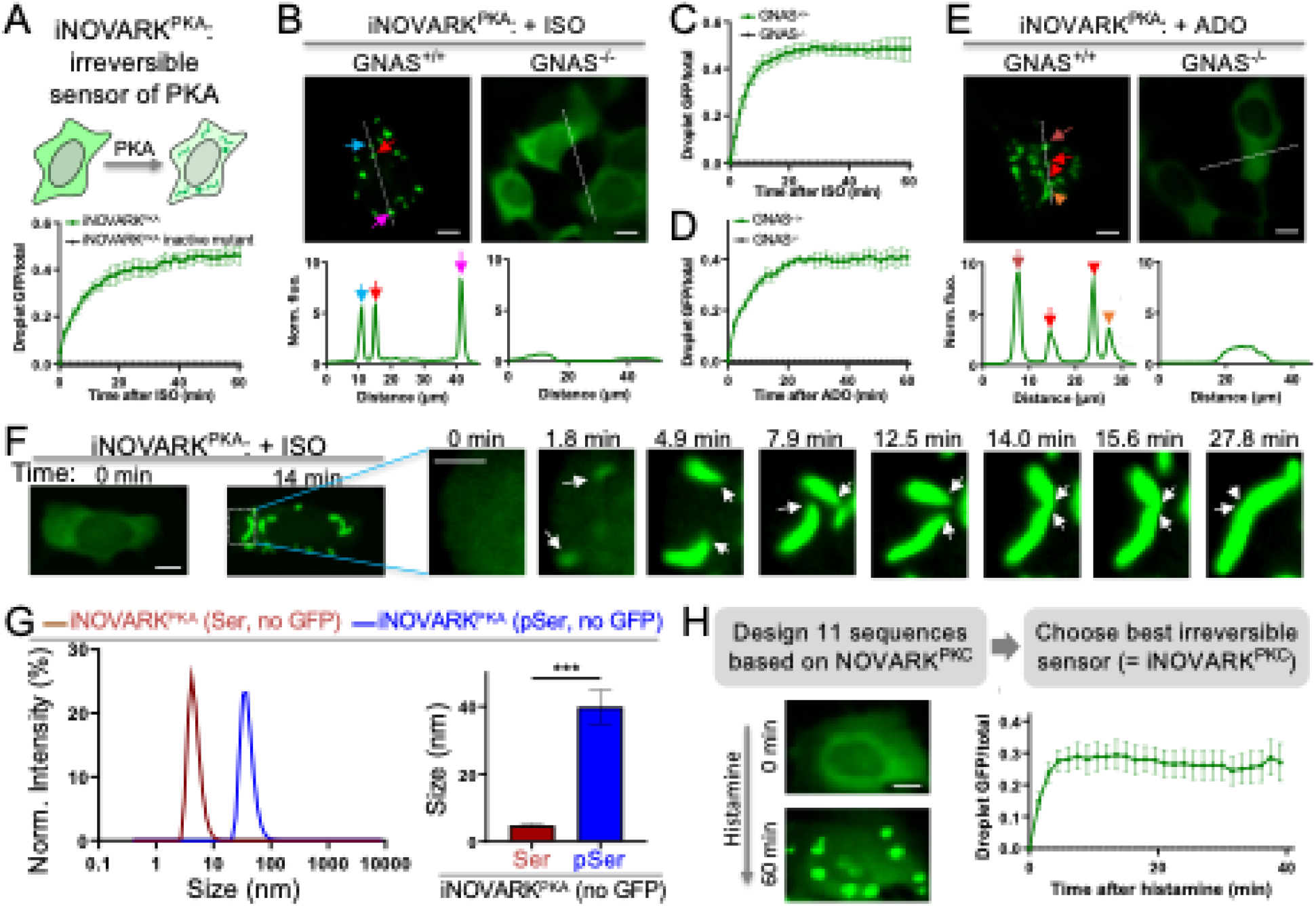
*De novo*-design of irreversible kinase biosensors. (A) Ratio of GFP in the higher-order assembly over total GFP as a function of time upon addition of ISO for iNOVARK^PKA^ and its inactive mutant that cannot be phosphorylated. Data are mean ± SE (n = 10 cells). (B – E) Fluorescence images and quantitative analysis of parent and GNAS knock-out HEK293 cells expressing iNOVARK^PKA^ upon addition of ISO (B, C) or adenosine (D, E). Histograms are shown along the dotted line in the images. Data are mean ± SE (n = 10 cells for C and D). (F) Time-lapse images showing formation of fibril-like structures via fusion of smaller structures.(G)Dynamic light scattering analysis of the phosphorylation-switch peptides in iNOVARK^PKA^. Data are mean ± SE (n = 6). (H) Fluorescence images and quantitative analysis of irreversible PKC biosensor iNOVARK^PKC^ upon addition of histamine. Data are mean ± SE (n = 10 cells). Scale bar, 10 μm (B, E, F, H); 5 μm (inset in F).

We next synthesized phosphorylated and non-phosphorylated peptide of the iNOVARK^PKA^. CD spectroscopy showed that both peptides did form helical structures (fig. S10 – 11). Dynamic light scattering showed that the phosphorylated peptide formed larger particles than the un-phosphorylated peptide (Fig. 4G, fig. S12), which is also larger than the reversible phospho-switch peptide (Fig. 1E).

To design an irreversible PKC reporter, we engineered the reversible reporter NOVARK^PKC^. We tested 11 mutants and found that upon histamine activation, one of them showed sustained signal that lasted for 40 minutes (Fig. 4H, table S2), in contrast to less than 6 minutes of the reversible reporter (Fig. 3G, H). The rest of the mutants showed no response, high background, or transient response (fig. S13).

Lastly, we examined whether the phospho-switch peptide is generalizable from green to other colors, such as red and near-infrared fluorescent proteins (FPs) (*34, 35*). Here, we used the irreversible PKA reporter as an example, and replaced GFP with other color FPs and found that the red FP mKO3 (*36*) and the near-infrared FP iRFP (*37*) are the best among them in forming higher-order assembly upon ISO (fig. S14). Some FPs such as mIFP (*38*) were also able to respond to the ISO-induced PKA activation, while others showed no response (fig. S15). These results indicate that for engineering multicolor *de novo*-designed reporters, different color FPs will need to be tested for each kinase reporters.

### Generalization of NOVARK to design reporters of a kinase that requires a docking site

While many kinases such as PKA and PKC phosphorylate their substrates by binding to the short sequence of the phospho-peptide, some kinases require an additional binding motif in order to bind and phosphorylate the substrates. One example is ERK that requires a docking motif for substrate phosphorylation. Here we decided to demonstrate that our *de novo*-designed NOVARK can incorporate such docking motifs and thus generalizable to engineer reporters for such kinases. First, we replaced the PKA substrate sequence by the ERK substrate sequence in the Design D (table S1 and 3). Then, we included the ERK docking motif to the N or C-terminal regions of the phospho-switch peptides. We tested 8 candidates. To induce ERK activity, we used EGF that binds to its receptor EGFR and activates ERK via the MAPK pathway (Fig. 5A). One of the candidates with the docking site at the C-terminal region showed response to EGF, while the rest showed no response or high background (fig. S16). However, the response of this candidate was slow, with the puncta formation requiring at least 14 minutes. Therefore, we decided to conduct a second round of engineering by optimizing this candidate. We tested 19 mutants and found that one of them showed strong response and the fluorescent puncta formed in ∼ 3 minutes upon addition of EGF (Fig. 5B – D). Furthermore, fluorescent puncta dissembled over time, indicating that the signal dissipated over time. This is consistent with transient ERK activation by EGF (*39*). We also demonstrated that puncta formation is dependent on the phospho-serine because mutation of this residue abolished the reporter signal (Fig. 5D, fig. S17). We further verified that the reporter response is specific to ERK activity by pretreating the cells with ERK inhibitor (ERKi) PD0325901, which abolished EGF-induced puncta formation (Fig. 5E – F).

**Fig. 5.**
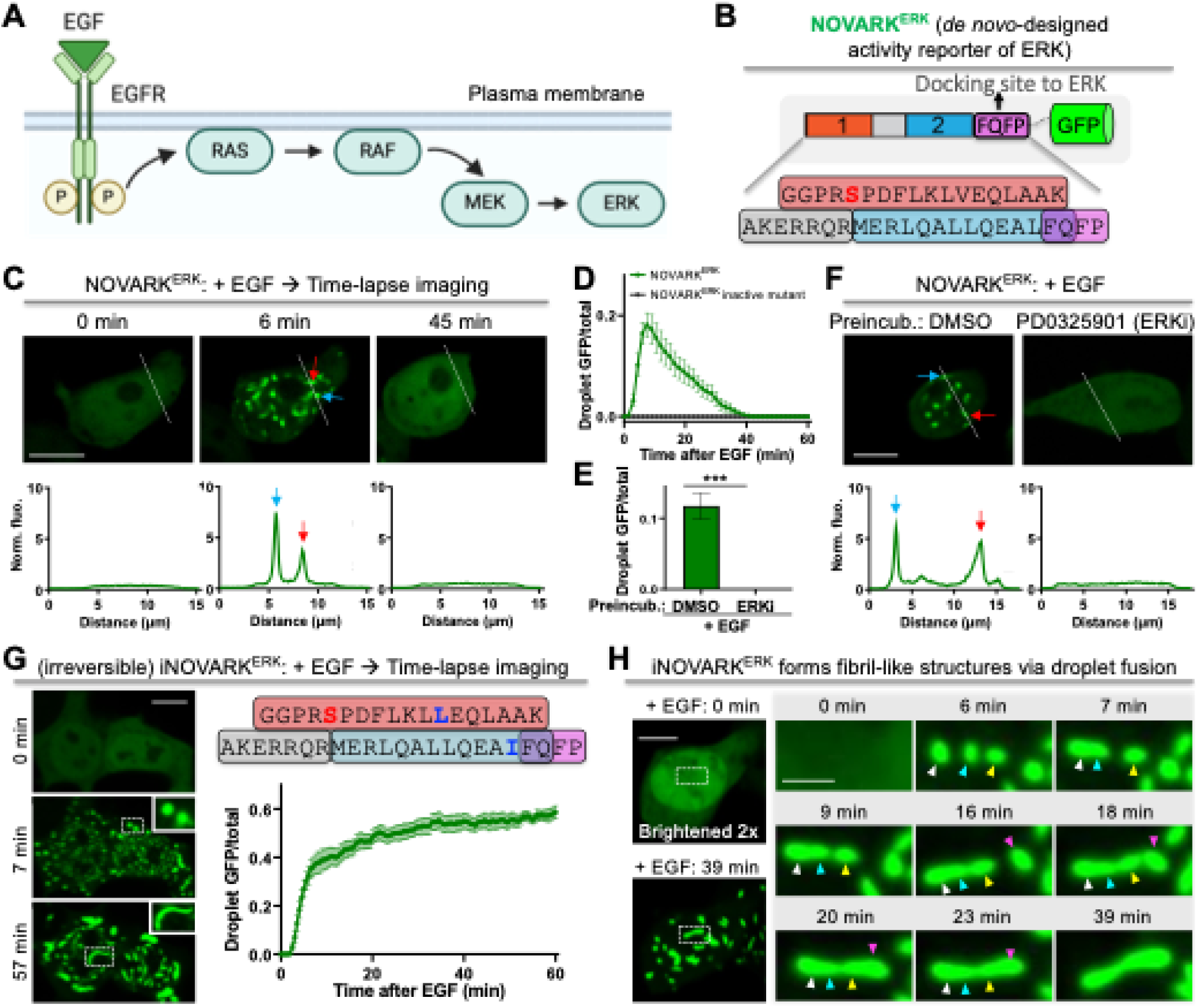
The NOVARK design can incorporate additional docking motifs for engineering ERK biosensor. (A) Schematic showing EGF-induced ERK activation. (B) Constructs of the ERK biosensor NOVARK^ERK^. The sequence of the phospho-switch peptide of the sensor is shown. The docking motif (FQFP) is shown in purple. Phospho-serine is shown in red. (C – F) Fluorescence images and quantitative analysis of HEK293T cells upon addition of EGF for NOVARK^ERK^ (C, D) and its inactive mutant that cannot be phosphorylated (D), and for NOVARK^ERK^ in cells that were pretreated with DMSO or the ERK inhibitor (ERKi) PD0325901 (E, F). Data are mean ± SE (n = 10 cells for D and E). (G) Fluorescence images and quantitative analysis of HEK293T cells expressing irreversible ERK biosensor iNOVARK^ERK^ upon addition of EGF. The sequence of the phospho-switch peptide of this sensor is shown on top-right. The two introduced mutations are shown in blue. Data are mean ± SE (n = 10 cells). (H) Time-lapse images showing formation of fibril-like structures via droplet fusion. Scale bar, 10 μm (C, F, G, H); 3 μm (inset in H).

Among the 19 mutants, we also found an irreversible ERK activity reporter that showed sustained signal in response to EGF. The higher-order assembly lasted for an hour upon EGF-induced ERK activation (Fig. 5G), in contrast to less than 40 minutes for a reversible ERK reporter (Fig. 5C, D). We named this reporter as iNOVARK^ERK^. Interestingly, this reporter formed round droplets first upon EGF (Fig. 5G, middle-left panel) and fibril-like structures later (Fig. 5G, bottom-left panel). Time-lapse imaging revealed that multiple round droplets fused together to form elongated fibril-like structures (Fig. 5H, movie S4). While iNOVARK^ERK^ was derived from Design D, it did not respond to ISO (fig. S18), consistent with our substrate change from PKA to ERK and the addition of ERK docking site. The rest of the 19 candidates showed slow response, weak and transient response, high background, or no response to EGF (fig. S19).

## Discussion

We sought to design a generalizable and modular kinase reporter system that is ultrabright with rapid kinetics, tunable signal duration, and fine spatial resolution. Our designed NOVARK system satisfies these requirements and is encoded by a peptide (∼40 residues) that is much smaller than existing reporter technologies. Importantly, we achieved these goals by testing limited number (10 to 20) of candidates with 1 to 2 rounds, and demonstrated that this reporter system is generalizable to multiple kinases. First, using biophysical first principles, we design phospho-switch peptides that self-assemble through symmetric, periodic interactions. Second, we select about 10 candidates and fuse them to GFP and test them in cells by examining whether they form higher-order assembly upon kinase activation. This combined approach of rational design and small-scale screening resulted in kinase reporters with large dynamic range, high brightness, and rapid kinetics with single-digit minute time-scale. Thus, our NOVARK reporters are orders of magnitude better than the previously reported *de novo*-based kinase reporters that suffer from slow kinetics requiring several hours to detect fluorescent signal (*8*). Our reporters are also advantageous than many FRET-based kinase reporters with orders of magnitude larger dynamic range and higher brightness (*28*). Furthermore, NOVARK reporters require no phosphopeptide-binding domain (PBD) that is necessary in many kinase reporters based on FRET or phase separation mechanisms (*12*-*14*). Exogenous expression of PBD might affect cell signaling because PBD may bind to the endogenous substrates in cells. Finally, the NOVARK reporters provided fine spatial resolution as demonstrated by NOVARK^PKC^ puncta originating at the membrane when the PKC is activated and recruited to the membrane by membrane-located stimulus PMA.

The NOVARK kinase reporters are demonstrated to be useful biological tools for studying signaling pathways, including the GPCR and RTK signaling that are two major signaling pathways in regulating cell growth and proliferation, and animal development, homeostasis and a diverse set of physiological functions (*40, 41*). The NOVARK biosensors are able to detect endogenous kinase activity with biologically relevant temporal dynamics for cell signaling activated via biological stimuli. In particular, the PKA reporter reads PKA signaling that is activated via G*α*s-coupled GPCR, including the *β*_2_AR and adenosine receptor upon addition of their agonist. The PKC biosensor reports PKC activity via G*α*q-coupled GPCR such as histamine receptor upon addition of histamine. Therefore, NOVARK will find important applications in studying GPCR signaling pathways in the future. Furthermore, we also showed that our design can incorporate a kinase docking site so that we can design reporters for kinases such as ERK that requires a docking site for substrate phosphorylation. And the ERK sensor reports EGF-induced endogenous ERK activity in living cells. Thus, our reporter will also find important applications in studying RTK signaling pathways that converge on ERK, including RTK regulators.

Our approach is generalizable to design reporters for other kinases, including those that require a docking site to the kinase. This paves the way for designing many other kinases. While reversible kinase reporters have advantage of monitoring dynamic activity of kinases, irreversible kinase reporters also have certain advantage using snapshot imaging, which will find applications in certain areas when time-lapse imaging is not preferred, such as high throughput screening of kinase inhibitors. Furthermore, our approach is compatible with multicolor kinase reporters by swapping out GFP with other FPs such as red and near-infrared FPs, though this requires further testing of the specific FPs since not all FPs are compatible with NOVARK.

In summary, we have developed a *de novo* design-based, generalizable approach for engineering kinase reporters, which paves the way to design most if not all 500 kinases in the future. These reporters have large dynamic range, high brightness and fast kinetics, and will find important applications in studying signaling pathways including but not limited to GPCR and RTK signaling. Our approach may also inspire biosensor design for other post-translational modifications such as methylation, acetylation, and ubiquitination.

## Supporting information

Supplementary Materials for Design of an ultrabright biosensor for dynamic imaging of kinase activity in cells

## Acknowledgments

Figures were generated with Biorender.com. We thank Drs. A. Inoue and D. Ma for sharing the cells, E. Q. Chen for generating the 3D movie, and Dr. R. Irannejad for constructive comments and suggestions. **Funding:** This research was support by: National Institute of Health R35GM131766 (to X.S.), R35GM122603 (to W.F.D), and F32GM147962 and K99GM155611 (to A.K.H). **Authors contributions:** Conceptualization: X.S., W.F.D.; Funding acquisition: W.F.D., X.S.; Investigation: all authors; Project Administration: X.S.; Supervision: W.F.D., X.S.; Visualization: X.L., S.K.T., C-I.C.; Writing – original draft: X.L., S.K.T., C-I. C., W.F.D., X.S.; Writing – review & editing: all authors. **Competing interests:** The authors declare the following competing interests: X.S., W.F.D., X.L., S.K.T. (University of California, San Francisco) have filed a patent application on the kinase biosensors. **Data and materials availability:** All data are available in the manuscript or the supplementary materials. The biosensor plasmid will be deposited to Addgene (https://www.addgene.org/).

## Supplementary Materials

Materials and Methods Figs. S1 to S19

Tables S1 to S4

References (*42*–*54*) Movies S1 to S4

