## Supplementary Materials for Design of an ultrabright biosensor for dynamic imaging of kinase activity in cells for "Design of an ultrabright biosensor for dynamic imaging of kinase activity in cells"

Materials and Methods

Figs. S1 to S19

Table S1 to S4

References

Movies S1 to S4

**Materials and Methods**

Cell culture

The 293T/17, HEK293, HEK293^GNAS-/-^ and Hela cells were passaged in Dulbecco’s Modified Eagle medium (DMEM) supplemented with 10% Fetal Bovine Serum (FBS), non-essential amino acids, penicillin (100 units/mL) and streptomycin (100 μg/mL). All culture supplies were obtained from the UCSF Cell Culture Facility.

Coiled-coil backbone parameterization

To achieve a self-assembling staggered topology, we generated idealized antiparallel 4-helix bundles using the CCCP program (*1*). The parameters were sampled around antiparallel 4-helix bundles found in nature: 3 values (6.5Å, 7.0Å, 7.5Å) were sampled for the superhelical radius parameter, 7 values (-4.4°, -4.1°, -3.8°, -3.5°, - 3.2°, -2.9°, and -2.6°) were sampled for the superhelical frequency parameter, 10 values (0° to 360° with 36° intervals) were sampled for the chain-wise alpha-helical phase parameter, and 5 values (-2.5Å, -1.25Å, 0Å, 1.25Å, 2.5Å) were sampled for the Z offset chain-wise parameter. In total, we generated 3*7*10*5 = 1,050 scaffolds.

Next, to determine the most viable scaffolds, we used MASTER (*2*) to survey the non-redundant PDB. We extracted a layer of 14 residues from the CCCP-generated coiled-coils (14 residues on each chain) and searched for matches in MASTER that met a 1Å RMSD criterion. Scaffolds that had at least 30 matches were determined to be sufficiently designable, and we proceeded with the 35 scaffolds that passed that filter.

To achieve the staggered effect, we removed every 6^th^ helical heptad (Fig. S1). Because the chains are antiparallel, the 6^th^ heptad voids are separated by two heptads where there is complete overlap between the 4 chains. Then, we grafted the 51 N-terminal phosphoserine-bearing motifs that we curated in our previous work (*3*) onto the N-termini of each isolated chain in our scaffold, sampling phosphoserines at the Ncap, N1, and N2 helical positions. An important principle guiding our design is that phosphoserines on the N-terminus of helical peptides (in our case, the kinase substrate) stabilize a helical conformation (*4*, *5*). We used structural bioinformatics to confirm this phenomenon in a previous study (*3*), in which we surveyed the PDB for phosphoserines at N-termini of helices. We found that majority of these phosphoserines adopt an N-capping rotamer, which allows their phosphate groups to hydrogen bond to unpaired backbone amides that are residing on the N-termini (fig. S2).

We stitched on the RRXS substrate motif of PKA, selected structures whose phosphoserines made at least two hydrogen bonds to its upstream or downstream backbone amides, selected for structures whose phosphoserine’s distal oxygens are close enough to an adjacent helix to potentially make an interhelical interaction (within 8Å), and eliminated structures where the phosphoserine clashed with the backbones of adjacent helices. This process whittled 35 scaffolds $\times$ 51 phosphoserine motifs $\times$ 3 N- terminal helical positions = 5,355 scaffolds down to 782 structures.

Sequence design

The 782 structures were then designed using ProteinMPNN (*6*). ProteinMPNN doesn’t recognize post-translationally modified amino acids, so to represent phosphoserine, we used glutamate, which is similarly charged and isosteric to phosphoserine. Sequences were symmetrized between chains by linking equivalent positions using a provided helper function that allows users to define homo-oligomers. Additionally, an arbitrarily high bias was placed on the substrate motif sequence to prevent mutation of those positions, and cysteines were completely omitted from sampling. We had to scan temperatures much higher than conventional sampling temperatures, because ProteinMPNN, in independent runs, designed long stretches of poly-alanine to practically the whole sequence, and we had to coax them out of their poly-alanine energy minima. We tried sampling temperatures of 0.25, 0.5, 0.75, 1, 2, and 5. ProteinMPNN generated 50 sequences for each sampling temperature, resulting in a total of 300 sequences per scaffold, for a grand total of 300 * 782 = 234,600 sequences.

To determine the most viable sequences, we ranked the sequences based on global score, percentage of alanines (< 30%), percentage of hydrophobic residues (30- 70%), and net charge (within -4 and 4, inclusive). We then took the top 1,000 sequences and used AlphaFold2 (*7*) to predict their structures, again using glutamate to represent phosphoserine. We evaluated 4-chain predictions of these sequences, because we observed that when we tried to predict higher-order oligomers, the structures optimized for packing and folded our sequences into globular proteins, rather than Extended, fibrillar coiled-coils. We then filtered the predicted structures to select for structures in which at least 2 of the 4 glutamate-representing phosphoserines form intermolecular salt bridges with an adjacent monomer, and selected for structures that were predicted to form anti-parallel, rather than parallel, coiled-coils. Lastly, we selected for structures that formed a staggered topology. An overwhelming majority of the designs were predicted to fold into 4-helix bundles with undesired topologies, with very few were predicted to stagger. The most common off-target topology featured the peptides aligning perpendicular to the superhelical axis, causing their termini to line up without the designed stagger (**fig. S1**). Another common result was that some predicted structures exhibited parallel topologies, rather than antiparallel. Therefore, the AlphaFold2 predictions had significant distinguishing power in isolating the best designs. The final sequences and models to be experimentally tested are listed in Table 1.

We designed 8 peptides containing the R-R-X-S substrate motif (DesignA, DesignB, DesignC, DesignD, Extended-DesignA, Extended-DesignB, Extended-DesignC and Extended-DesignD, **Table S1**). Designs A-D were directly derived from sequence design, and each of their “Extended” counterparts were created by inserting an additional repeat of a corresponding internal heptads to strengthen coiled-coil association (**Table S1**).

Plasmid construction

All plasmid constructs were created by standard molecular biology techniques and confirmed by exhaustively sequencing the cloned fragments. Each reporter was constructed with a substrate-*de novo* peptide (see **Table S1**) and GFP sequence, or red and near-infrared fluorescent proteins

(*8*-*13*). To create each reporter candidates, DNA fragment with most consensus sequence and BsmBI flanked variable region were inserted into pcDNA3 plasmid upstream of GFP sequence to generate a shuttle vector, cut with BsmBI then a short DNA fragment, encoding variable region and with complementary overhang for shuttle vector, was synthesized and annealed to ligate into the shuttle vector.

Live cell imaging

Cells were grown on Nunc^®^ Lab-Tek^®^ II chambered coverglass for imaging experiment.

293T/17 and HEK293 cells were transiently transfected using calcium phosphate transfection reagent with 100 ng plasmid. Hela cells were transiently transfected using Lipofectamine 3000 Transfection Reagent with 100 ng. Imaging was carried out 1 day after transfection. For drug pretreatment, cells were preincubated in respective inhibitor for around 1 hour. Concentration of the small molecules was used as: forskolin 1 µM, Isoproterenol 10 µM, H89 10 µM, Histamine 50 µM, PMA 1 µM, GO6893 10 µM, EGF 10 ng/ml and PD0325901 10 µM. Imaging was carried out on Nikon Eclipse Ti inverted microscope equipped with Yokogawa CSU-W1 confocal scanner unit (Andor), digital CMOS camera ORCA-Flash4.0 (Hamamatsu) and ASI MS-2000 XYZ automated stage (Applied Scientific Instrumentation). Imaging was performed in environmental control unit incubation chamber (*In Vivo* Scientific) maintained at 37 °C and with 5% CO_2_. Fluorescence images were acquired using Nikon CFI Plan Apochromatic 20X dry (N.A. 0.75) objective or CFI apochromatic TIRF 60X oil objective (N.A. 1.49) against GFP, Cherry and IFP *per* experiment settings.

Solid phase peptide synthesis and purification

*De novo* designed peptides were synthesized using an Fmoc-protection strategy via solid phase peptide synthesis with an Initiator+ Alstra peptide synthesizer (Biotage) on Tentagel S-RAM resin. Phosphoserine was installed synthetically by incorporating Fmoc-Ser(PO(OBzl)OH-OH in place of Fmoc-Ser-OH. Peptides were acetylated with HCTU/DIPEA/AcOH prior to cleavage from resin and global deprotection with a 95/2.5/2.5 TFA/TIPS/H2O cleavage cocktail. Cleaved peptides were precipitated in cold ether before purification via high-performance liquid chromatography (HPLC). Peptide mass was confirmed by MALDI-TOF mass spectrometry and purity confirmed by analytical HPLC. HPLC-purified peptides were lyophilized before use in in vitro experiments.

*In vitro* biophysical characterization of *de novo* peptides

Circular dichroism (CD): Lyophilized peptides were dissolved in buffer (10 mM MOPS, pH 7.5). Protein concentration was measured by UV-Vis and calculated by absorbance at A280. Secondary structure was characterized by circular dichroism (Jasco J-810 spectropolarimeter). Spectra were collected from 190–250nm with 0.5nm data pitch in continuous scanning mode at 50 nm/min and 1 nm band width, and twelve accumulations were collected per sample. Thermal denaturation experiments were conducted with a temperature gradient from 20 to 90°C and ramp rate of 1°C/min with ellipticity at 222nm measured every 2°C. Thermal denaturation with chemical denaturant gradients were conducted with a 0-6M urea gradient.

Dynamic light scattering (DLS): Peptide samples at 100 and 200 µM in 10 mM MOPS, pH 7.5 were prepared for DLS. DLS measurements were collected using a Malvern Zetasizer Nano S90. Each sample was analyzed with 3 or 6 runs of 20 measurements each. Distribution means were calculated from the average measured size across all runs.

Transmission electron microscopy (TEM)

Peptide stock samples were prepared from lyophilized stocks in buffer (10 mM MOPS, pH 7.5, 100 mM NaCl) concentration was measured by UV-Vis. Samples for TEM were prepared to the final desired concentration from stocks. Negatively stain samples were prepared using 400-mesh carbon-coated copper grids (SPI). Briefly, an aliquot of sample (5 µL) was incubated for 30 seconds on a glow-discharged grid, and the solution was removed by filter paper blotting. Grids were washed 3x with ddH_2_O followed by staining 3x with 0.75% (w/v) uranyl formate (Electron Microscopy Sciences) with vacuum aspiration. Stained samples were imaged with a Talos L120C operated at 120 keV, and micrographs were recorded using a Ceta-D (Thermo Fisher Scientific) camera.

Data quantification and plotting

All images were processed in Image J with “Analyze Particle function” to score the sum of droplets pixel fluorescence intensity and the cells pixel intensity (Droplet GFP/total = GFP in the higher-order assembly over total GFP). All data were quantified and plotted using Graphpad Prism 10.

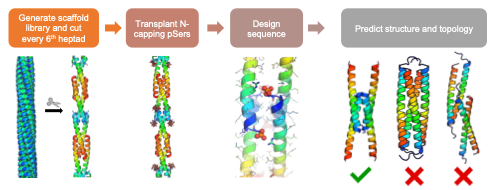

**Figure S1. The design pipeline starts with parameterizing symmetric 4-helix bundles, removing every 5th heptad to stagger each peptide, capping the N-termini with phosphoserine, designing the sequence with ProteinMPNN, and filtering on AlphaFold2 structure prediction.**

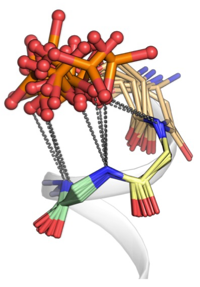

**Figure S2. Phosphoserines capping N-termini of alpha-helices.** Representative N-terminal helical phosphoserines, taken from crystallographically-solved structures in the PDB, demonstrate N-capping potential by hydrogen bonding to the exposed, unpaired backbone amides that are also residing on the N- terminus, thus stabilizing alpha-helical conformation.

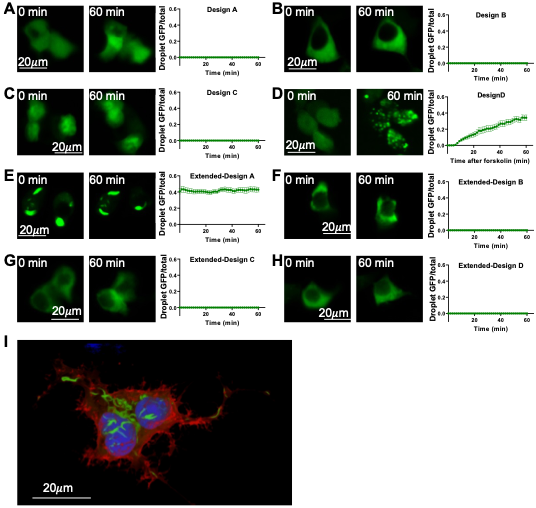

**Figure S3. Fluorescence imaging of de novo-designed PKA reporters. Note these reporters are named as Design A – D; and extended Design A – D.** (A to H) Representative images and quantitative analysis showing performance of 8 candidates (Design A – D, extended Design A – D, respectively) before and after forskolin stimulation. (I) 100X image shows fibril-like structure of Extended-Design A. Data are mean ± SE (n = 10 cells).

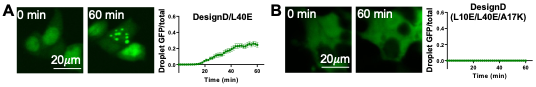

**Figure S4. Optimization based on Design D for PKA reporter.** (**A** and **B**) Representative images and quantitative analysis showing performance of other two candidates before and after forskolin stimulation. Data are mean ± SE (n = 10 cells).

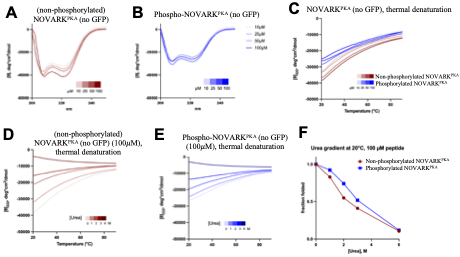

**Figure S5. *In vitro* characterization of NOVARK^PKA^. (A** and **B**). Concentration-dependent circular dichroism of and NOVARK^PKA^ ± phosphoserine show that all variants have helical secondary structure in *vitro*. **C**. Concentration-dependent thermal denaturation with monitoring of secondary structure via circular dichroism at 222nm for NOVARK^PKA^ ± phosphoserine. (**D, E and F)**. Thermal and chemical denaturation in the presence of varied [urea] with monitoring of secondary structure via circular dichroism at 222nm for NOVARK^PKA^ ± phosphoserine.

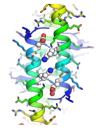

**Figure S6. Predicted structure of NOVARK^PKA^ + phosphoserine .**

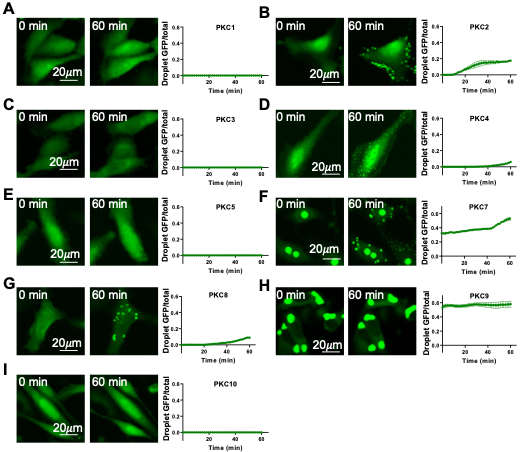

**Figure S7. Initial development of PKC reporters.** Here the PKC reporters are named as PKC1 – 10. Representative images and quantitative analysis showing performance of other nine candidates after PMA stimulation. Note: PKC6 is shown in the main figures and named as NOVARK^PKC^. Data are mean ± SE (n = 10 cells).

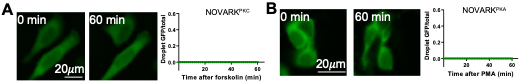

**Figure S8. Fluorescence imaging of NOVARK^PKC^ in response to forskolin stimulation (A) and NOVARK^PKA^ in response to PMA stimulation (B).** Data are mean ± SE (n = 10 cells).

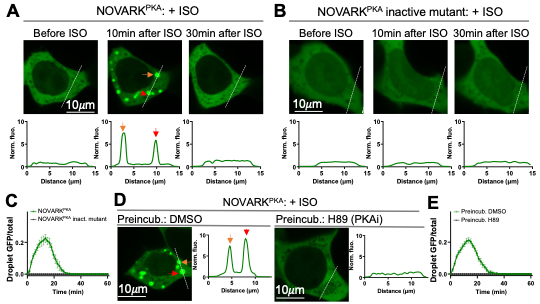

**Figure S9. Characterization of NOVARK^PKA^.** (**A**, **B**) Representative showing NOVARK^PKA^ (**A**) and the inactive mutant (Ser to Arg mutation) (**B**), the lower panels show fluorescence intensity over distance along the white line shown in the images. (**C**) Quantitative analysis before and after isoprenaline application. (**D, E**) Representative images (**D**) and quantitative analysis (**E**) showing performance of NOVARK^PKA^ after preincubation with DMSO or PKA inhibitor (H89). Data are mean ± SE (n = 10 cells).

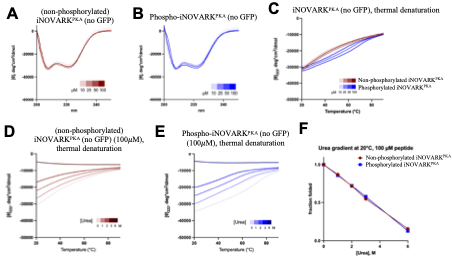

**Figure S10. *In vitro* characterization of iNOVARK^PKA^.** (**A** and **B**). Concentration-dependent circular dichroism of and iNOVARK^PKA^ ± phosphoserine shows that all variants have helical secondary structure *in vitro*. (**C**). Concentration-dependent thermal denaturation by monitoring the secondary structure via circular dichroism at 222nm for iNOVARK^PKA^ ± phosphoserine. (**D, E,** **F**). Thermal and chemical denaturation in the presence of varied [urea] by monitoring the secondary structure via circular dichroism at 222nm for iNOVARK^PKA^ ± phosphoserine.

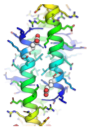

**Figure S11. Predicted structure of iNOVARK^PKA^ + phosphoserine.**

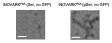

**Figure S12. Representative negative stain TEM micrographs of iNOVARK^PKA^ with unphosphorylated (left) and phosphorylated (right) serine.** Scale bar, 100 nm.

continued in next page

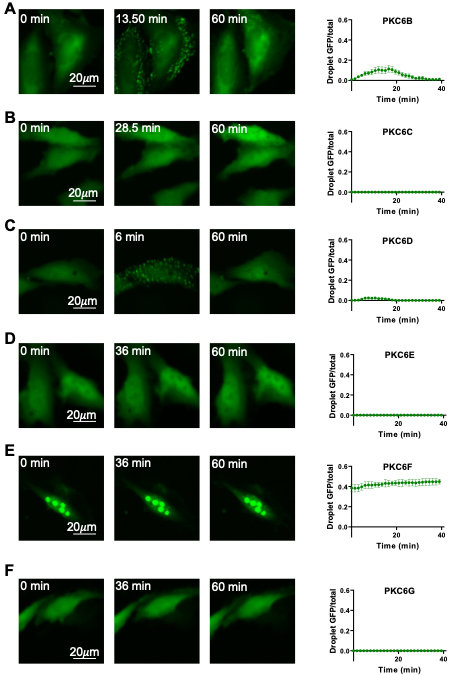

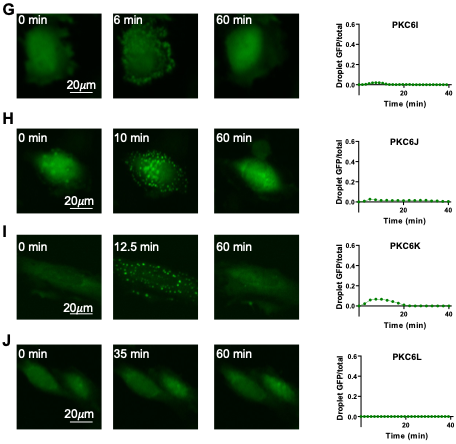

**Figure S13. Development of irreversible PKC reporters.** Here the PKC reporters are named as PKC6B – L. Representative images and quantitative analysis showing performance of these candidates after histamine stimulation. Data are mean ± SE (n = 10 cells).

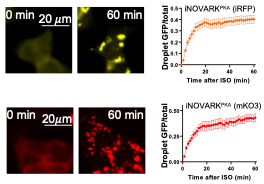

**Figure S14. Fluorescence images of the best near-infrared iNOVARK^PKA^ (iRFP) and red iNOVARK^PKA^ (mKO3) in response to isoprenaline stimulation.** Data are mean ± SE (n = 10 cells).

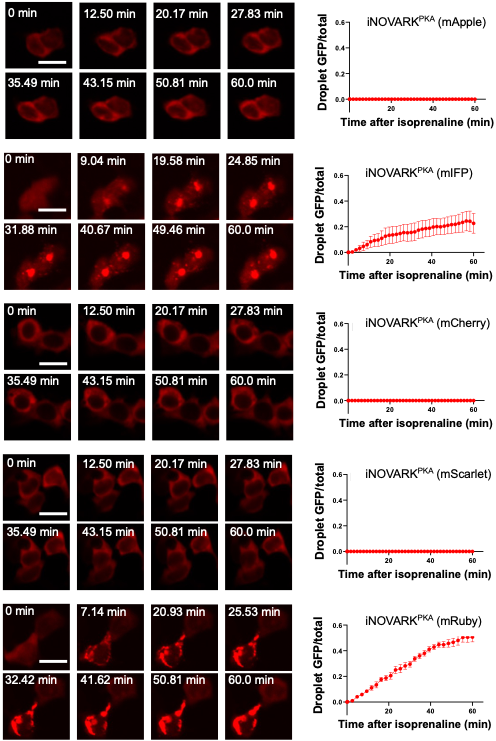

**Figure S15. Fluorescence imaging of iNOVARK^PKA^ using different fluorescent proteins in response to isoprenaline stimulation.** Data are mean ± SE (n = 10 cells). Scale bar, 20 μm.

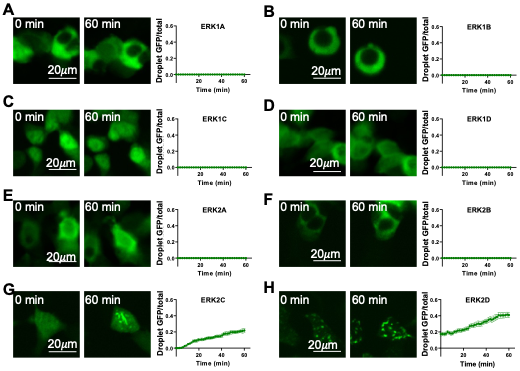

**Figure S16. Initial development of ERK reporters (the 1st round).** Here the ERK reporters are named as ERK1A – D; and ERK2A – D. Representative images and quantitative analysis showing performance of eight candidates after EGF stimulation from which ERK2C is selected for further optimization. Data are mean ± SE (n = 10 cells).

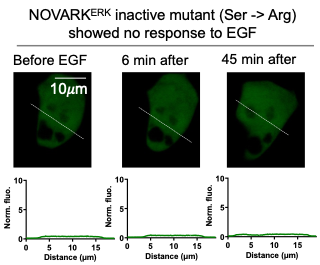

**Figure S17.** Representative images of the inactive mutant (Serine->Arginine) of NOVARK^ERK^ in response to EGF stimulation. The lower panels are the fluorescence intensity over distance along the white line in shown the images (upper panels).

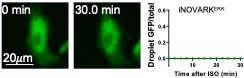

**Figure S18. Representative images and quantitative analysis showing iNOVARK^ERK^ in response to isoprenaline.** Data are mean ± SE (n = 10 cells).

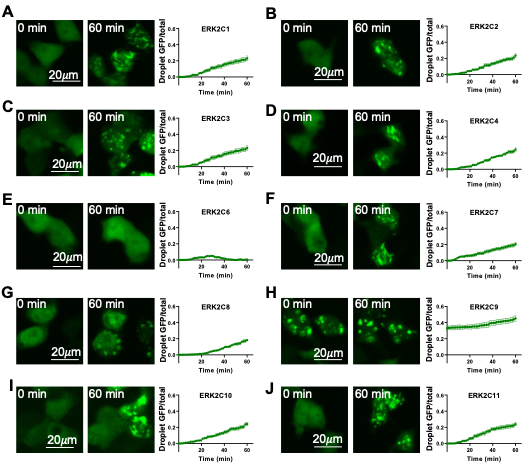

Continued in next page

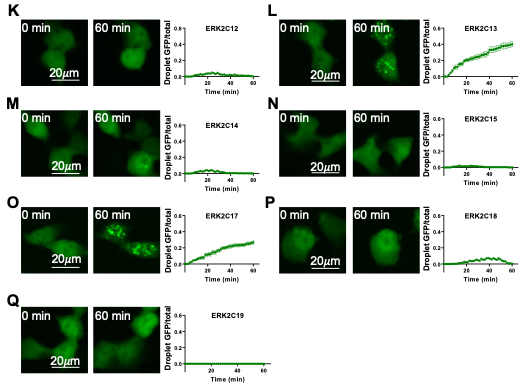

**Figure S19. Optimization of the ERK reporters (the 2nd round).** Here the ERK reporters are named as ERK2C1 – 19. Representative images and quantitative analysis showing performance of these candidates after EGF stimulation. Data are mean ± SE (n = 10 cells).

**Table S1. List of tested phosphorylation-switch peptides for sensing PKA activity.** The reporters were initially named as Design A – D, and extended Design A – D.

|  | **Peptide sequence (RRR(S) is the PKA substrate motif. (S) marks phosphorylation site)** |
| --- | --- |
| Design A | GGRRR(**S**)ILAKVKQAIEELEQRLQERQQDVENLEALTKRMR |
| Design B | GGRRR(**S**)LQVLKNLTQRAEAARQREEVLHQELDRARQG |
| Design C | GGRRR(**S**)LEALLALLARLQEFAEREARLRALLEEARRL |
| **Design D** | GGRRR(**S**)LEFLKLLEQLAAKAKERRQRMERLQALLQEALRL |
| Extended-Design A | GGRRR(**S**)ILAKVKQAIEELEQREQEAQQDVENLEALTKRMRARA |
| Extended-Design B | GGRRR(**S**)LQVLKNLTQRAEAARQREEVLHQELDRARQAQAA |
| Extended-Design C | GGRRR(**S**)LEALLALLARLQERAEALARLSALLEEARRLLARE |
| Extended-Design D | GGRRR(**S**)LEFLKLLEQLAAKAKERRQRMERLQALLQEALRLAAAL |
| **Design D**/L40E | GGRRR(**S**)LEFLKLLEQLAAKAKERRQRMERLQALLQEALRE |
| **Design D**/ L10E/A17K/L40E | GGRRR(**S**)LEFEKLLEQLKAKAKERRQRMERLQALLQEALRE |
| **Design D**/L10E/A17K (NOVARK^PKA^) | GGRRR(**S**)LEFEKLLEQLKAKAKERRQRMERLQALLQEALRL |
| **Design D**/L10E (iNOVARK^PKA^) | GGRRR(**S**)LEFEKLLEQLAAKAKERRQRMERLQALLQEALRL |
| NOVARK^PKA^ inactive mutant (S to R) | GGRRR(**R**)LEFEKLLEQLKAKAKERRQRMERLQALLQEALRL |
| iNOVARK^PKA^ inactive mutant (S to R) | GGRRR(**R**)LEFEKLLEQLAAKAKERRQRMERLQALLQEALRL |

**Table S2. List of tested phosphorylation-switch peptides for sensing PKC activity.** The reporters were initially named as PKC1 – 10; and PKC6A – L.

|  | **Peptide sequence (RFRRFQS is the PKC substrate motif. (S) marks phosphorylation site)** |
| --- | --- |
| PKC1 | GGRFRRFQ(**S**)LKDKALLEQLAAKAKERRQRMERLQALLQEALRL |
| PKC2 | GGRFRRFQ(**S**)LKDFALLEQLAAKAKERRQRMERLQALLQEALRL |
| PKC3 | GGRFRRFQ(**S**)LKDKALLEQLAAKAEERRQRMERLQALLQEALRL |
| PKC4 | GGRFRRFQ(**S**)LKFFALLEQLAAKAKERRQRMERLQALLQEALRL |
| PKC5 | GGRFRRFQ(**S**)LKFKALLEQLKAKAKERRQRMERLQALLQEALRL |
| PKC6 (NOVARK^PKC^) | GGRFRRFQ(**S**)LKDFALLEQLAAKAEERRQRMERLQALLQEALRL |
| SR-PKC6 | GGRFRRFQ(**R**)LKDFALLEQLAAKAEERRQRMERLQALLQEALRL |
| PKC7 | GGRFRRFQ(**S**)LKFFALLEQLAAEAKERRQRMERLQALLQEALRL |
| PKC8 | GGRFRRFQ(**S**)LKFFALLEQLAAKAEERRQRMERLQALLQEALRL |
| PKC9 | GGRFRRFQ(**S**)LKDFALLEQLAAEAEERRQRMERLQALLQEALRL |
| PKC10 | GGRFRRFQ(**S**)LKFFALLEQLKAEAKERRQRMERLQALLQEALRL |
| PKC6A (iNOVARK^PKC^) | GGRFRRFQ(**S**)LKDEALLEQLAAKAEERRQRMERLQALLQEALRL |
| PKC6B | GGRFRRFQ(**S**)LKDFAALEQLAAKAEERRQRMERLQALLQEALRL |
| PKC6C | GGRFRRFQ(**S**)LKDFALLEQLAAKAEERRQRMERLQAALQEALRL |
| PKC6D | GGRFRRFQ(**S**)LKDFALLEQLAAKAEERRQRMERLQALLQEAIRL |
| PKC6E | GGRFRRFQ(**S**)LKDFALLEQLAAKAEERRQRMERLQALLQEALRL |
| PKC6F | GGRFRRFQ(**S**)LKFFALLEQLAAKAEERRQRMERLQALLQEALRL |
| PKC6G | GGRFRRFQ(**S**)LKDFALLEQLKAKAEERRQRMERLQALLQEALRL |
| PKC6I | GGRFRRFQ(**S**)LKDFALLEQLAAKAKERRQRMERLQALLQEALRL |
| PKC6J | GGRFRRKQ(**S**)LKDFALLEQLAAKAEERRQRMERLQALLQEALRL |
| PKC6K | GGRFRRFG(**S**)LKDFALLEQLAAKAEERRQRMERLQALLQEALRL |
| PKC6L | GGRFRRFQ(**S**)LKKFALLEQLAAKAEERRQRMERLQALLQEALRL |

**Table S3. List of tested phosphorylation-switch peptides for sensing ERK activity.** The reporters were initially named as ERK1A – D, ERK2A - D; and ERK2C1 – 19.

|  | **Peptide sequence (PR(S) is the ERK substrate motif. (S) marks phosphorylation site. FQFP is the docking site to ERK)** |
| --- | --- |
| ERK1A | GGPR(**S**)PAFLKLLEQLAAKAKERRQRMERLQALLQEFQFP |
| ERK1B | GGPR(**S**)PAFLKLLEQLAAKAKERRQRMERLQALLQEAFQFP |
| ERK1C | GGPR(**S**)PAFLKLLEQLAAKAKERRQRMERLQALLQEALFQFP |
| ERK1D | FQFPGGSAGGSAGGSAGGSAGGPR(**S**)PAFLKLLEQLAAKAKER RQRMERLQALLQEALRL |
| ERK2A | GGPR(**S**)PDFLKLLEQLAAKAKERRQRMERLQALLQEFQFP |
| ERK2B | GGPR(**S**)PDFLKLLEQLAAKAKERRQRMERLQALLQEAFQFP |
| ERK2C | GGPR(**S**)PDFLKLLEQLAAKAKERRQRMERLQALLQEALFQFP |
| ERK2D | FQFPGGSAGGSAGGSAGGSAGGPR(**S**)PDFLKLLEQLAAKAKER RQRMERLQALLQEALRL |
| ERK2C1 | GGPR(**S**)PDFLKILEQLAAKAKERRQRMERLQALLQEALFQFP |
| ERK2C2 | GGPR(**S**)PDFLKVLEQLAAKAKERRQRMERLQALLQEALFQFP |
| ERK2C3 | GGPR(**S**)PDFLKALEQLAAKAKERRQRMERLQALLQEALFQFP |
| ERK2C4 | GGPR(**S**)PDFLKLIEQLAAKAKERRQRMERLQALLQEALFQFP |
| ERK2C5 (NOVARK^ERK^) | GGPR(**S**)PDFLKLVEQLAAKAKERRQRMERLQALLQEALFQFP |
| SR-ERK2C5 | GGPR(**R**)PDFLKLVEQLAAKAKERRQRMERLQALLQEALFQFP |
| ERK2C6 | GGPR(**S**)PDFLKLLEQIAAKAKERRQRMERLQALLQEALFQFP |
| ERK2C7 | GGPR(**S**)PDFLKLLEQVAAKAKERRQRMERLQALLQEALFQFP |
| ERK2C8 | GGPR(**S**)PDFLKLLEQLAAKAKERRQRMERIQALLQEALFQFP |
| ERK2C9 | GGPR(**S**)PDFLKLLEQLAAKAKERRQRMERVQALLQEALFQFP |
| ERK2C10 | GGPR(**S**)PDFLKLLEQLAAKAKERRQRMERAQALLQEALFQFP |
| ERK2C11 | GGPR(**S**)PDFLKLLEQLAAKAKERRQRMERLQAILQEALFQFP |
| ERK2C12 | GGPR(**S**)PDFLKLLEQLAAKAKERRQRMERLQAVLQEALFQFP |
| ERK2C13 | GGPR(**S**)PDFLKLLEQLAAKAKERRQRMERLQAALQEALFQFP |
| ERK2C14 | GGPR(**S**)PDFLKLLEQLAAKAKERRQRMERLQALIQEALFQFP |
| ERK2C15 | GGPR(**S**)PDFLKLLEQLAAKAKERRQRMERLQALVQEALFQFP |
| ERK2C16 (iNOVARK^ERK^) | GGPR(**S**)PDFLKLLEQLAAKAKERRQRMERLQALLQEAIFQFP |
| ERK2C17 | GGPR(**S**)PDFLKLLEQLAAKAKERRQRMERLQALLQEAVFQFP |
| ERK2C18 | GGPR(**S**)PDFLKLLEQLAAKAKERRQRMERLQALLQEAAFQFP |
| ERK2C19 | GGPR(**S**)PEFLKLLEQLAAKAKERRQRMERLQALLQEALFQFP |

**Table S4. List of reagents and cell lines in the study.**

| **REAGENTS** | **SOURCE** | **IDENTIFIER** |
| --- | --- | --- |
| Chemicals, Peptides, and Recombinant Proteins | | |
| Calcium Phosphate Transfection Kit | ThermoFisher Scientific | K278001 |
| Lipofectamine 3000 Transfection | ThermoFisher Scientific | L3000001 |
| Forskolin | TargetMol | T2939 |
| Isoproterenol | Sigma-Aldrich | 1351005; CAS: 51-30-9 |
| Adenosine | Sigma-Aldrich | A9251; CAS: 58-61-7 |
| Phorbol-12-myristate-13-acetate, PMA | Sigma-Aldrich | 524400 |
| Histamine dihydrochloride | TargetMol | T6534 |
| Epidermal Growth Factor human, EGF | Sigma-Aldrich | E9644 |
| PKA inhibitor, H-89 dihydrochloride hydrate | Sigma-Aldrich | B1427; CAS:130964-39-5 (anhydrous) |
| PKC inhibitor, GO6893 | Sigma-Aldrich | G1918 |
| ERK inhibitor, PD 0325901 | Sigma-Aldrich | PZ0162; CAS: 391210-10-9 |
| Experimental Models: Cell Lines | | |
| 293T/17 | ATCC | CRL-11268 |
| Hela | Dengke Ma, UCSF | N/A |
| (Parent) HEK293 cells | A. Inoue (Tohoku University) | N/A |
| HEK293^GNAS-/-^ cells | A. Inoue (Tohoku University) | N/A |
| Recombinant DNA | | |
| Plasmid: pcDNA3-PKA reporters | This paper | N/A |
| Plasmid: pcDNA3- PKA reporters | This paper | N/A |
| Plasmid: pcDNA3-ERK reporters | This paper | N/A |
| Plasmid: pmIFP-H2B-T2A-CAAX-mCherry | This paper | N/A |
| Software and Algorithms | | |
| NIS-Element | Nikon | N/A |
| Fiji ImageJ | Opensource | https://fiji.sc/ |

**Movie S1. 3D images of Extended-Design A-GFP in a HEK293T cell.** The designed peptide (extended-design-D) fused to GFP is shown in green. The membrane targeted mCherry-CAAX is shown in red. The nucleus-targeted H1B-mIFP is shown in blue**.**

**Movie S2. NOVARK^PKC^ reports membrane-associated PKC activation after PMA stimulation.** NOVARK^PKC^ and mCherry-CAAX were co-transfected into HeLa cells. PMA was added to induce PKC activation. The membrane targeted mCherry-CAAX is shown in red**.**

**Movie S3. Response of NOVARK^PKC^ after histamine stimulation.** NOVARK^PKC^ and mCherry-CAAX were co-transfected into HeLa cells. Histamine was added to induce PKC activation. The membrane targeted mCherry-CAAX is shown in red**.**

**Movie S4. Fusion event of iNOVARK^ERK^ after EGF stimulation.** iNOVARK^ERK^ was transfected into HEK293T cells. EGF was added to induce ERK activation.
